# Infection with gut parasites correlates with gut microbiome diversity across human populations in Africa

**DOI:** 10.1101/2025.01.23.634469

**Authors:** Mirabeau M. Ngwese, Bayode R. Adegbite, Jeannot F. Zinsou, J. Liam Fitzstevens, Victor T. Schmidt, Alvyn N. Moure, Moustapha M. Maloum, Alexander V. Tyakht, Kelsey E. Huus, Nicholas D. Youngblut, Peter G. Kremsner, Ayola A. Adegnika, Ruth E. Ley

## Abstract

Soil-transmitted helminths (STH) are common in (sub)tropical regions and primarily affect impoverished populations. STHs reside in the gut, interacting both with the gut microbiota and host immunity. Clinical STH detection is laborious and often not performed within the context of gut microbiome studies. Here, we assessed whether fecal metagenome data could be used to assess STH infection, and to relate STH infection to microbiome diversity. We generated 310 gut metagenomes for mother-child pairs from two different locations in Gabon: one rural and one semi-urban. The presence and abundance of four STH parasites (*Ascaris lumbricoides*, *Strongyloides stercoralis*, *Trichuris trichiura*, and *Necator americanus*) were assessed using both microscopy and qPCR. Sequence data were used to characterize the microbiomes and to detect the presence of these four STH parasites. We found that metagenome data could accurately detect the presence of all four STHs, as reflected by high sensitivity and specificity compared to microscopy or qPCR detection. Furthermore, the number of STH species present in stool was associated with microbiome diversity and with the abundances of specific taxa, most notably in young children. We applied this approach to microbiome data from five other populations in Africa and corroborated our findings. Our results demonstrate that human intestinal parasites can be accurately detected via metagenomic sequencing, and highlight how infection with multiple STH parasites is linked to consistent features in the gut microbiome in populations in Africa.

## Introduction

Soil-transmitted helminth infections are endemic in developing countries, with over 1.5 billion people - representing 24% of the global population - affected by soil-transmitted helminths (STH) ^1^. These infections are predominantly found in tropical and subtropical regions ^2^, particularly sub-Saharan Africa ^2,3^. Transmission occurs through oral-fecal contact via contaminated soil and transcutaneously through larval penetration of the skin. Infections can spread within and between communities through shared environments. The prevalence of STHs in rural communities is often poorly documented, especially in countries with limited resources and lacking mass drug administration (MDA) programs. MDA strategies, including deworming of high-risk populations and the widespread administration of anthelmintic drugs, are essential for controlling and reducing the morbidity associated with worm infections ^4^.

The World Health Organization (WHO) classifies the STH species *Ascaris lumbricoides*, *Trichuris trichiura*, Hookworms (*Necator americanus* and *Ancylostoma duodenale)* and *Strongyloides stercoralis* as neglected tropical diseases. These first three STHs contributed to an estimated 1.9 million disability-adjusted life years worldwide in 2019 ^5–7^. Strongyloidiasis, caused by *S. stercoralis*, further affects an estimated 30 to 100 million people worldwide ^8^. At-risk groups include young children, preschool and school-age children, adolescent girls, women of reproductive age, pregnant women and adults with certain high-risk jobs such as tea-pickers or miners ^5,9^. STH parasites often cause significant nutritional deficiencies, affecting growth and cognitive development in children ^10^. They can also impact the behavior of the patient and lead to intestinal inflammation and malabsorption ^11–13^.

STHs are members of the gut microbiome. As such, they interact with other components of the microbiome such as Bacteria and Archaea, and these interactions can impact the success of parasite colonization and the outcome of parasitic infections ^14^. Conversely, parasite colonization can affect the host’s interaction with the microbiome ^15–17^. When parasites colonize the gut, changes occur in the gut barrier, such as the epithelial mucus layers, which can subsequently alter the composition of microbiota ^18,19^. Helminths can influence host-microbiome interactions by modulating immune signaling pathways and altering nutrient availability, thereby creating conditions that support their persistence within the gut ecosystem ^20^. Furthermore, helminth-induced immunomodulation fosters a tolerogenic environment, which may encourage the growth of specific microbial taxa, indicating a potential mutualistic relationship between helminths and certain microbial communities ^21^. Therefore, interactions between parasites and microbes may play significant roles in modifying host physiology and influencing disease outcomes ^22^.

The investigation of parasite-microbiome relationships across diverse populations has been limited in part because quantifying STH load is laborious and time-intensive and is not always done in conjunction with microbiome studies. Recently, a number of groups have reported the possibility of detecting parasite presence by metagenomic sequencing in livestock stool ^23,24^ and the detection of unicellular parasites, such as Blastocystis, in human fecal metagenomes ^25^. Although metagenomic sequencing shows promise for detecting a wide range of pathogens including STH, its diagnostic performance has not yet been fully validated against traditional diagnostic methods such as microscopy and PCR ^26^, highlighting the need for further standardization and targeted validation studies.

Here, we examined STH load in subjects sampled in Gabon using conventional techniques (microscopy, qPCR) to compare with estimates generated from gut metagenomes. We then examined the relationship between STH infection with gut microbiome diversity from the same individuals. Subjects consisted of mother-child pairs from two different communities: the semi-urban region of Lambaréné, where parasite prevalence is well-documented ^27,28^, and the rural "Ikobey" region, which lacks parasite distribution data. We then expanded our analysis by leveraging publicly available metagenomic data from five additional studies in diverse African populations. Our results indicate that metagenome data can be used to assess STH richness, revealing reproducible patterns in microbiome-parasite associations across populations.

## Results

### Prevalence of parasites in samples from Gabon

We collected 310 human stool samples from mothers and their children in Gabon (Figure 1A) described in ^29^. This dataset comprises matched mother-child pairs from two distinct regions, the rural Ikobey (n=40 pairs) and the semi-urban Lambaréné (n=115 pairs). The average age of adults was 27 ± 6.7 (s.d.) years, while that of children was 40 ± 45 (s.d.) weeks. Adults and children tended on average to be older in Ikobey compared to Lambaréné (Wilcoxon rank sum test, p = 6e-04 and p = 0.0032, respectively; Table 1). We focused on the four STH species previously reported as prevalent in this population *(i.e.*, *A. lumbricoides*, *T. trichiura*, *N. americanus* and *S. stercoralis*) ^30,31^. We processed 222 of the 230 Lambaréné stool samples for STH via microscopy analysis for eggs/larvae at the Centre de Recherches Médicales de Lambaréné (CERMEL) in Gabon. All samples (310) were then shipped to Germany for metagenome sequencing generated in ^29^, and for qPCR detection of STHs, which included 230 samples from Lambaréné and 80 from Ikobey.

**Figure 1:**
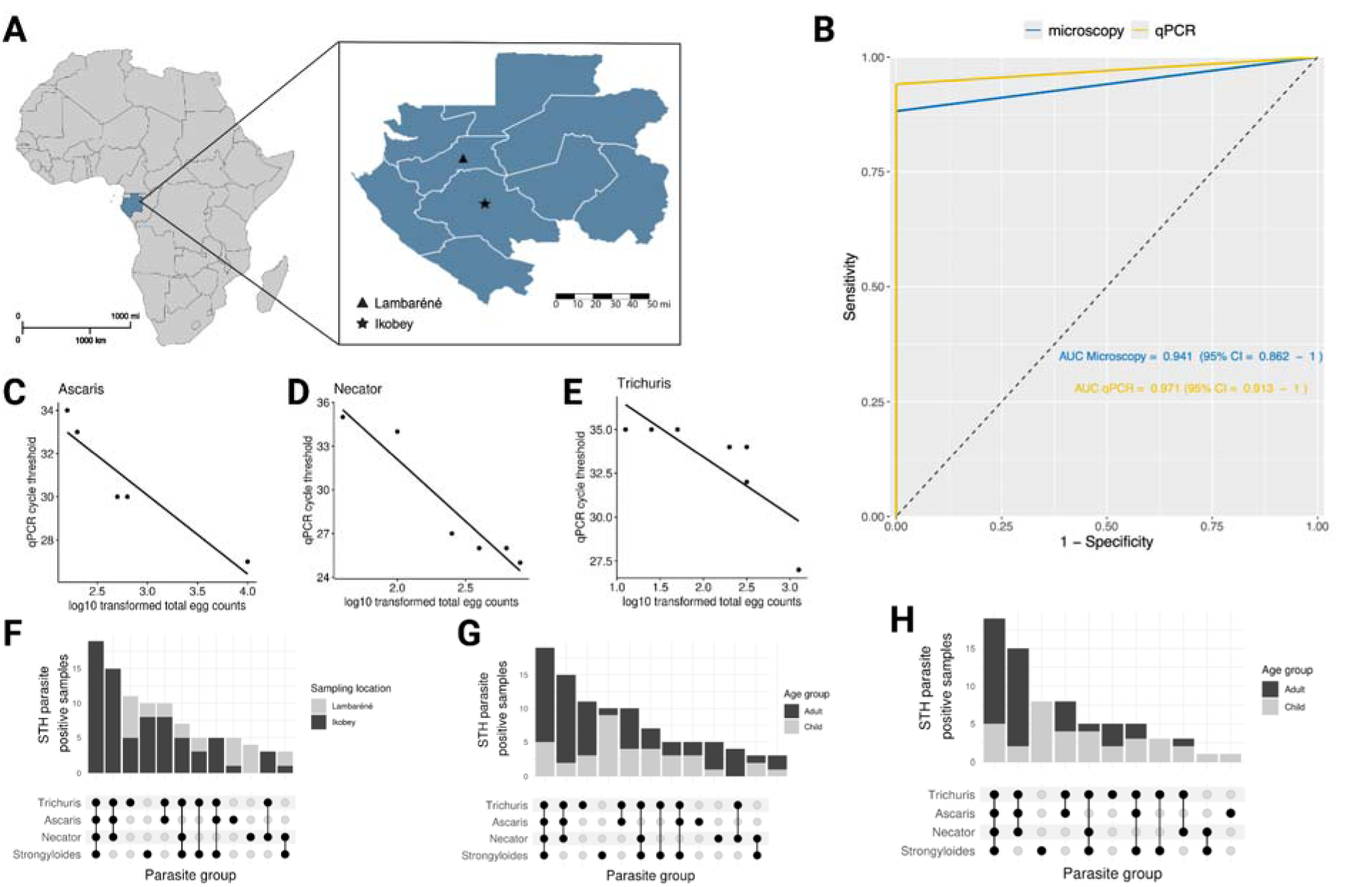
Comparison of microscopy and qPCR performance for detecting soil-transmitted helminths (STHs) and assessing the prevalence of four STH species in Lambaréné and Ikobey. **A)** Map of Gabon showing the sampling locations (= Ikobey villages, ▲ = Lambaréné). **B)** Area under the ROC curve (AUC) comparing the performance of microscopy and qPCR for detecting STH parasites. **C-E**. Correlation between microscopy-based egg counts (per gram of stool) and qPCR-based abundance (Ct values) for STH species: **C)** Correlation between qPCR cycle threshold values (y-axis) and log10-transformed total egg counts per gram of stool for *Ascaris lumbricoides* (*R =* 0.97*, p* = 0.0048). **D)** Correlation for *Necator americanus* (*R =* 0.93*, p* = 0.002). **E)** Correlation for *Trichuris trichiura* (*R =* 0.99*, p* = 0.00031). **F)** Total number of STH-positive samples based on combined microscopy and qPCR data, displayed by sampling location. **G)** Total number of STH-positive samples from combined microscopy and qPCR data, displayed by age group. **H)** Total number of STH-positive samples from Ikobey, determined by qPCR data only.

Among the 222 Lambaréné samples processed by microscopy, 14 tested positive for STH, and all of these were also confirmed by qPCR, indicating a high overlap between the two methods, with qPCR detecting additional cases not identified by microscopy. To assess the accuracy of both methods for detecting STH parasites, we defined a “true positive” as any sample that tested positive by either microscopy, qPCR, or both. Overall, qPCR demonstrated higher sensitivity (94%, CI: 0.713–0.999) and specificity (100%, CI: 0.983–1.000) compared to microscopy, which showed a sensitivity of 88% (CI: 0.636–0.985) and specificity of 100% (CI: 0.983–1.000). To assess the ability of both methods to distinguish between samples with and without STH parasites, and to evaluate their agreement in quantifying individual parasites, we performed ROC curve and correlation analyses (Figure 1B-E). The AUC values were 0.941 (95% CI: 0.862–1.000) for microscopy and 0.971 (95% CI: 0.913–1.000) for qPCR, demonstrating similar overall performance (Figure 1B). DeLong’s test showed no significant difference between the two AUCs (p = 0.5716). We observed strong correlations (absolute Spearman correlation > 0.9, p < 0.005) between qPCR cycle threshold values and log10-transformed total egg counts per gram of stool for *A. lumbricoides*, *N. americanus*, and *T. trichiura* (Figure 1C-E). Based on this analysis, we proceeded to analyze the Ikobey samples using qPCR only, and observed infection rates of 95% in adults and 88.5% in children (Table 1).

To gain an overview of the parasite load in adults and children in both locations, we combined the Lambaréné and Ikobey results. We observed all four parasites in both locations (Figure 1F) and across both adult and child age groups (Figure 1G). However, parasite infections were significantly less common in children compared to adults (Table 1 and Table S1a-c), with the majority of infections occurring in the rural area of Ikobey (Figure 1H). Additionally, infection patterns differed by location: rural Ikobey exhibited a higher prevalence of parasite infections and a greater number of parasite species per sample on average (Figure 1H), compared to the semi-urban area of Lambaréné, regardless of age group (Table 1 and Table S1d).

### Associations of gut microbiome alpha and beta diversity with number of STH species

We next sought to investigate associations between the microbiome and parasite load estimated by qPCR (n=310). To evaluate gut microbiome diversity at the species level, we employed metagenomic profiling against a custom database created from the Genome Taxonomy Database (GTDB Release 202) ^32^. This approach significantly improved the classification of reads compared to the standard Kraken database, increasing the percentage of classified reads from 38.0% ± 26.4 (s.d.) to 81.8% ± 5.6 (s.d.). Using generalized linear models, we investigated the relationship between microbiome and parasite for both measures of alpha diversity (species counts or richness), while accounting for location and age group. As noted above, the parasite alpha diversity (species count) differed significantly between adults and children. For both children and adults however, we observed that samples with a higher parasite species count tended to have a richer microbiome (measured with Shannon diversity index and Faith’s PD; Figure 2A and B, Tables S2a and S2b).

**Figure 2:**
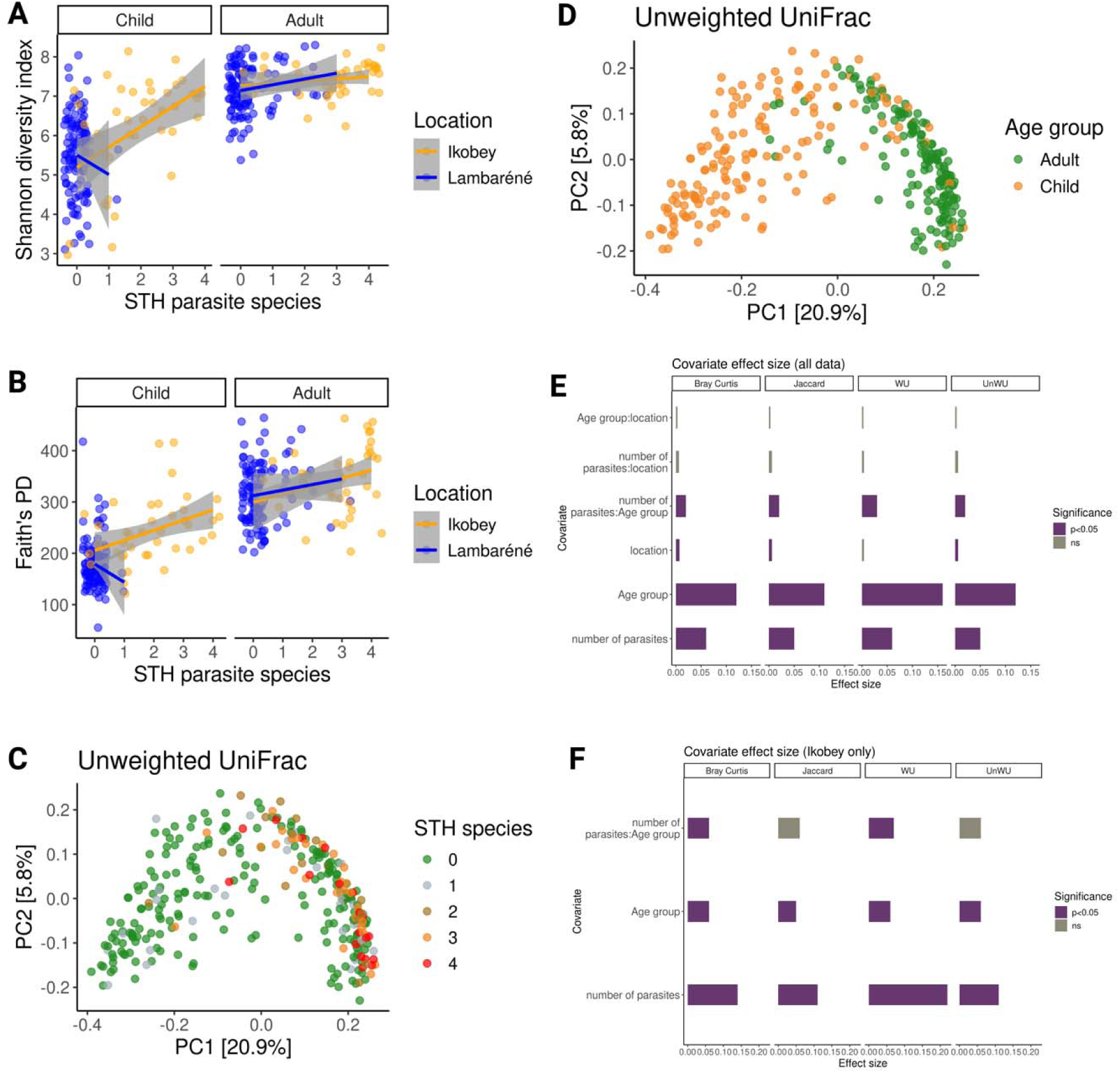
Gut Microbiome Diversity in Relation to STH species count, Age Group, and Location. **A)** Correlation between gut microbiome Shannon diversity index and STH species count for adults and children by location (correlation method: GLM). For children, *R^2^* = 0.23 in Ikobey and *R^2^* < 0.01 in Lambaréné; for adults, *R^2^* = 0.02 in both Ikobey and Lambaréné. **B)** Correlation between Faith’s Phylogenetic Diversity (PD) and STH species count for adults and children by location (correlation method: GLM). For children, *R^2^* = 0.15 in Ikobey and *R^2^* = 0.01 in Lambaréné; for adults, *R^2^* = 0.07 in Ikobey and *R^2^* = 0.01 in Lambaréné. **C)** Principal Coordinates Analysis (PCoA) of unweighted UniFrac values for all samples (n = 310), colored by STH species count. **D)** PCoA of unweighted UniFrac values for all samples, colored by age group. **E)** Effect size (R²) of each covariate plotted against significance (p < 0.05 = *, ns = non-significant) for all samples. **F)** Effect size (R²) of each covariate plotted against significance for Ikobey samples only. Effect sizes were calculated using four beta diversity metrics: Bray-Curtis, Jaccard, Weighted UniFrac, and Unweighted UniFrac. Abbreviations: WU = Weighted UniFrac, UnWU = Unweighted UniFrac.

To better account for the impact of age on the relationship between microbiome and parasite diversity, we focused on samples with available age information (n = 307). In children, as expected, age was strongly associated with microbiome richness; however, microbiome richness remained significantly associated with parasite species count after accounting for age (p = 4.55e-05, p = 3.48e-09, respectively; Table S2c), whereas no significant associations were observed in adults (n = 155; Table S2d, Figure S1A). Faith’s PD was not associated with parasite species count in either age group but was significantly associated with age in children, with no such association in adults (Table S2e & Table S2f, Figure S1B). These findings suggest that in children, both the richness and evenness of the microbiome are important in explaining the diversity of STH parasite species, and this relationship is influenced by age. However, in adults, these associations are weaker, indicating that age plays a critical role in shaping the microbiome-parasite relationship, with the microbiome potentially becoming more stable and less influenced by parasites as individuals age.

We then investigated the between-sample diversity (beta diversity) of the microbiome in relation to age and the parasite species count. We observed that the beta-diversity of the microbiome was strongly influenced by age group, and was also influenced by location, as visualized by Principal Coordinate Analysis (PCoA) (Figure 2C&D) and tested via PERMANOVA (Figure 2E & F, Figure S1C, Table S3a). Furthermore, the parasite species-count had a significant, albeit modest, association with microbiome beta diversity (Figure 2E, Figure S1D) when controlling for both location and age (PERMANOVA test: Bray-Curtis: *p* = 0.001, Jaccard: *p* = 0.001, Weighted UniFrac: *p* = 0.001, Unweighted UniFrac: *p* = 0.001, Table S3a). This effect was particularly pronounced in the Ikobey population (n=80), where parasite prevalence was highest (Figure 2F, Table S3b). The effect size of STH species count on beta diversity was also greatest when using Weighted UniFrac distance (Figure 2F), suggesting that the number of parasites better explains differences in the abundances of most abundant taxa. In summary, the parasite count in two regions of Gabon was strongly associated with gut microbiome alpha and beta diversity.

### Specific microbial taxa are differentially abundant by STH number of species

Having observed significant differences in microbiome diversity by parasite-richness, we next sought to identify specific microbial taxa correlating with parasite richness. To account for the substantial differences in microbiome composition between age groups, we conducted separate analyses for adults and children. To ensure robustness, we employed two linear modeling approaches, DESeq2 and MaAslin2, which utilize different normalization methods ^33,34^.For adults, DESeq2 identified 106 differentially abundant species associated with the number of parasite species, while MaAslin2 identified 26 (padj < 0.05, Table S3c). Only 14 species (12%) were identified by both methods. These overlapping species are shown in Figure 3A, and their relative abundances, considering only these 14 species, are displayed in Figure 3B. Among children, DESeq2 identified 77 species and MaAslin2 identified 98, with 39 overlapping species (29%) (Table S3d). The relative abundances of the overlapping species are shown in Figure 3C, and the log2 fold changes are displayed in Figure 3D. No species overlapped between adults and children.

**Figure 3:**
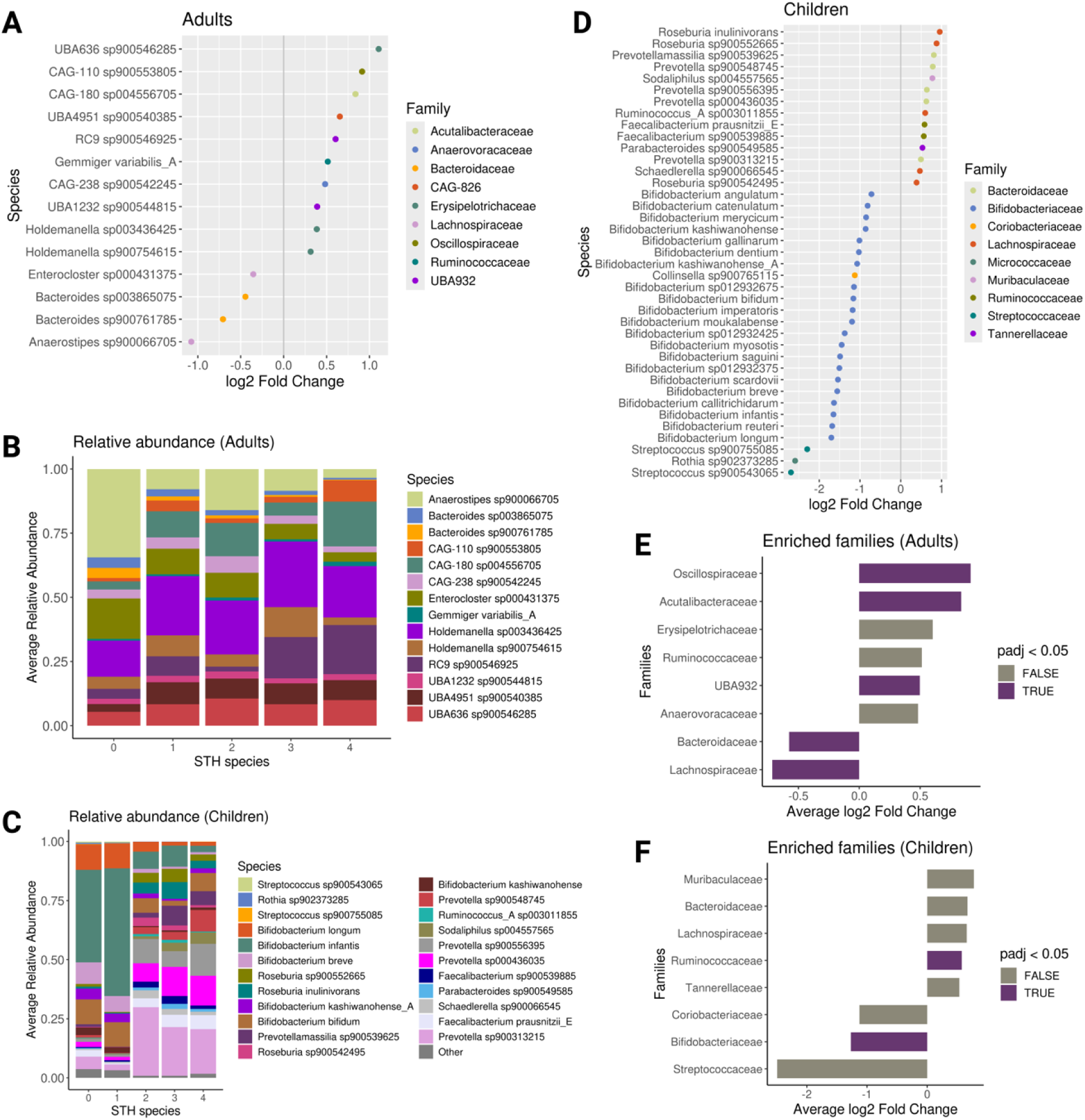
Differentially abundant microbial species by STH parasite species count, with no overlap between adults and children. **A)** Log2 fold changes in microbial species abundance associated with STH parasite count in adults, determined by DESeq2 and MaAslin2, adjusting for location. Only species identified as significant by both methods (Adj. p < 0.05) are shown. **B)** Relative abundances of differentially abundant species overlapping between DESeq2 and MaAslin2 in adults. **C)** Relative abundances of differentially abundant species overlapping between DESeq2 and MaAslin2 in children. **D)** Log2 fold changes in microbial species abundance associated with STH parasite count in children, determined by DESeq2 and MaAslin2, adjusting for location. Only significant species (Adj. p < 0.05) identified by both methods are shown. **E)** Average log2 fold changes of bacterial families significantly enriched in parasite-negative samples in adults, based on Fast Gene Set Enrichment Analysis (FGSEA). **F)** Average log2 fold changes of bacterial families significantly enriched in parasite-negative samples in children, based on FGSEA. Microbial species abundances were pre-filtered to include only species with at least 80% prevalence across samples in both adults and children before performing the differential abundance analysis. The log2 fold changes are derived from DESeq2, with significance confirmed by both DESeq2 and MaAslin2.

Focusing on differentially abundant taxa identified as significant by both methods, we observed that among the 14 species in adults, 4 were negatively associated with parasite species count, representing the Bacteroidaceae (n = 2) and Lachnospiraceae (n = 2). The remaining 10 species were positively associated with parasites, encompassing 7 families: Erysipelotrichaceae, UBA932 (Bacteroidales), Ruminococcaceae, Oscillospiraceae, Anaerovoracaceae, Acutalibacteraceae, and CAG-826 (Bacilli). In children, out of the 39 differentially abundant species, 25 were negatively associated with the number of parasites, primarily belonging to the Bifidobacterium family (n = 21), followed by Streptococcaceae (n = 2), and one species each from Micrococcaceae and Coriobacteriaceae (Figure 3D). Using an algorithm commonly used for gene set enrichment analysis, we performed a fast gene set enrichment analysis (FGSEA) and identified taxonomic families that were significantly overrepresented among differentially abundant species in both adult and child datasets ^35^. Among adults, we found 5 families, primarily from the Firmicutes phylum (n = 3), that were significantly overrepresented among microbes positively (n = 3) or negatively (n = 2) associated with the number of parasites (padj<0.05; Figure 3E, Table S3e). In contrast, the children dataset revealed only 2 families, Ruminococcaceae and Bifidobacteriaceae, which were significantly enriched or depleted with the number of parasites among the differentially abundant species (padj <0.05; Figure 3F, Table S3f). These findings suggest that the relationship between parasite species counts and bacterial species is complex. While some bacterial species increase with higher parasite counts (positive association), others decrease (negative association). In adults, a stronger positive association is observed between parasite counts and various bacterial species. In contrast, many species in children, particularly from the Bifidobacterium family, show a negative association with parasite counts, suggesting that the presence of certain beneficial bacteria is lower in individuals with higher parasite counts. Additionally, certain families, such as Ruminococcaceae, play a key role, showing both positive and negative associations depending on the age group. Overall, the relationship between parasites and gut microbes varies by microbial family and host age.

### Quantification of parasites by qPCR correlates with metagenome sequence data

Next, we explored the feasibility of detecting and quantifying STH parasites directly from shotgun metagenome data. After filtering out bacterial and archaeal sequences, we employed KrakenUniq ^36^ to classify the remaining sequences using a custom database of the four STH parasites. KrakenUniq output provided k-mer counts mapped to each STH reference genome. All four STH species were reliably detected (presence/absence) by shotgun metagenomics (Figure S2A). Notably, this approach significantly improved on existing reference-based metagenomic methods such as EukDetect ^37^ which was not able to detect the prevalent human STHs A. *lumbricoides*, *T. trichiura*, or *N. americanus* (Figure S2B).

We evaluated the sensitivity and specificity of our method against our qPCR data. Out of the 310 samples, the metagenome STH parasite k-mer counts method identified 97 samples positive for at least one STH parasite and 213 negative samples. This represents a sensitivity of 82,47% and a specificity of 87,32%, indicating that this method is a reliable approach for detecting STH infections (Table S11).

We next assessed whether metagenome data could be used for within-sample quantification of parasite load. We observed strong and significant positive correlations between the abundances of three out of the four species inferred by qPCR and metagenomics (Figure 4 A-D; Spearman Rank Correlation: p < 3.4e-8; r > 0.66 for all 3 species). However, the counts for *T. trichiura* did not correlate between qPCR and metagenome-based methods (p = 0.12; r = 0.16). Our analysis revealed that while the qPCR primers for *A. lumbricoides, N. americanus* and *S. stercoralis* specifically matched their target species in Blast alignments, the reverse primer for *T. trichiura* had a stronger E-value of 0.047 for non-target nematodes, such as *Calodium hepaticum.* This cross reactivity with other nematodes can potentially cause inaccuracies in qPCR results.

**Figure 4:**
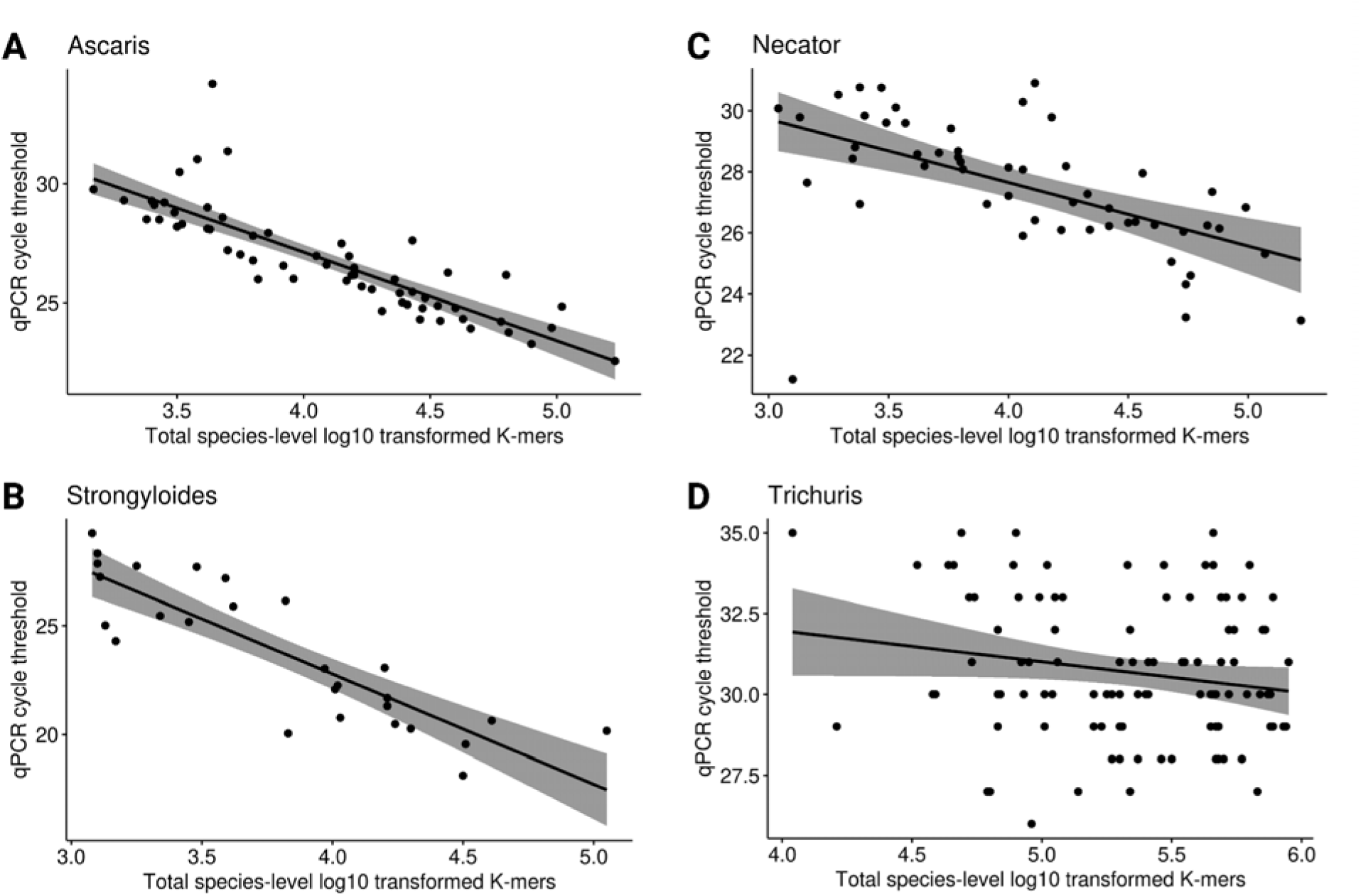
Correlation between metagenome-quantified STH parasite abundances and qPCR. **A)** Correlation between qPCR cycle threshold values and log10-transformed species-specific k-mer counts for *Ascaris lumbricoides* (*R =* 0.91*, p <* 2.22 X 10^-16^). **B)** Correlation between qPCR cycle threshold values and log10-transformed species-specific k-mer counts for *Strongyloides stercoralis* (*R =* 0.88*, p =* 1.8 X 10^-9^). **C)** Correlation between qPCR cycle threshold values and log10-transformed species-specific k-mer counts for *Necator americanus* (*R =* 0.67*, p =* 3.5 X 10^-8^). **D)** Correlation between qPCR cycle threshold values and log10-transformed species-specific k-mer counts for *Trichuris trichiura* (*R =* 0.16*, p =* 0.12). `Spearman’s rank-based correlation test was used to assess significance, with only correlations showing p-values ≤ 0.05 considered significant`.

Quantifying multicellular organisms from metagenomic data is inherently complex due to variability in genome size, gene copy number, and the presence of different life stages in the sample. Additionally, sample preparation and DNA extraction methods may differentially affect the recovery of *T. trichiura* DNA. These combined factors underscore the challenges in accurately quantifying *T. trichiura* using both qPCR and metagenomic read count data, as translation of gene abundances into total organism numbers is inherently difficult. Because of these challenges, we adopted a conservative approach that allows for the inclusion of more samples by using the presence or absence of STH species in our metagenomic data for subsequent analyses rather than the parasite load.

### Co-occurrence of the gut microbiome and STH parasites

Using probabilistic models based on metagenomic data, we inferred co-occurrences between microbes and STH parasites presence/absence. In adults, all 4 STH parasites showed co-occurrences with specific microbial species (Figure 5A&B), while in children, only 3 parasites (excluding Trichuris) exhibited such co-occurrences (Figure 5C&D). Consistent with the association between parasite species count and increased microbial diversity, parasite species presence tended to be positively associated with bacterial presence, especially in adults (Figure 5B&E). Ascaris had the highest number of unique co-occurrences with microbes in both adults and children (Figure 5E&F). Positive associations were frequently observed between parasites and Firmicutes or Bacteroidota, while negative associations were observed between parasites and Actinobacteriota (Figure 5E&F). Furthermore, analysis of random co-occurrence graph subsets indicated a low probability of associations between parasites or bacteria with similar properties (Figure S2C). Overall, our findings support the associations of microbial diversity with the number of parasites and suggest that these STH parasites may influence specific microbial niche compositions.

**Figure 5:**
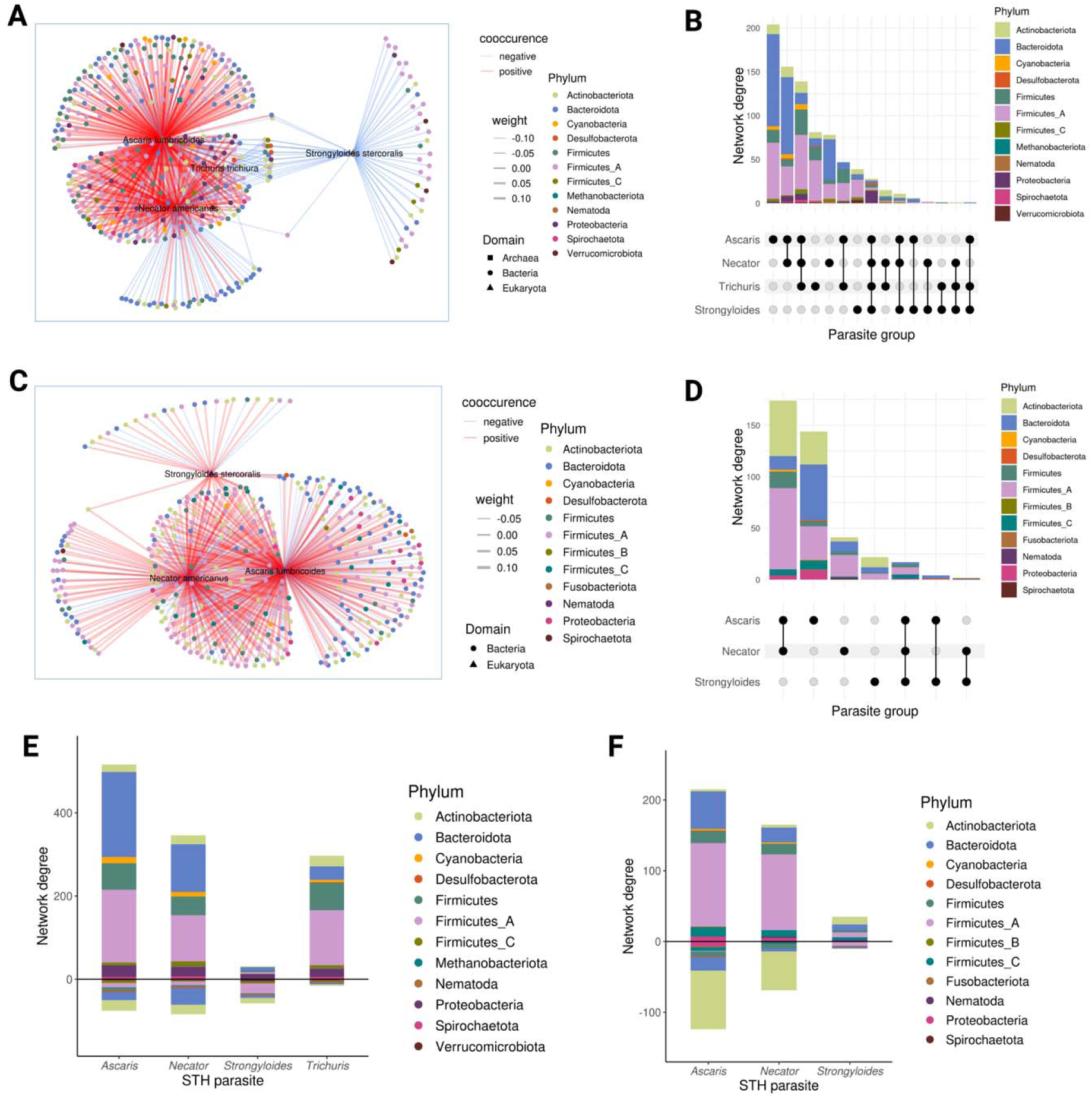
Co-occurrence network plots for adults and children in the Gabon dataset. **A)** Co-occurrence network for adults, displaying species with significant co-occurrence probabilities. **B)** Network degree (number of connections per node) within the adult co-occurrence network. **C)** Upset plot showing parasites linked to similar bacteria/archaea in the adult network. **D)** Co-occurrence network for children, displaying species with significant co-occurrence probabilities. **E)** Network degree (number of connections per node) within the children’s co-occurrence network. **F)** Upset plot showing parasites linked to similar bacteria/archaea in the children’s network. Nodes represent species with significant co-occurrence probabilities, with red indicating positive co-occurrence and blue indicating negative co-occurrence. Only species associations with weights between -0.14 and -0.01 (negative) and 0.05 to 0.15 (positive) are shown for visualization.

### Consistent parasite-microbiome associations across African datasets

Using metagenomic data to infer parasite species counts allowed us to leverage publicly available datasets for the analysis of parasite-microbiome associations. We conducted a meta-analysis to determine the generalizability of parasite-microbe associations in African populations using five metagenome datasets from Gabon (this study), Cameroon^25,38^, Burkina Faso ^39^, Ethiopia ^40^, and Madagascar ^41^, as well as an American reference cohort ^42^. Since metagenomic data was extremely limited for young children from these regions, the analysis focused on adult women (mean age = 32) to align with our previous analyses. A total of 466 (410 African samples) samples were included in the meta-analysis (Figure 6A). All four STH species were detected in the African datasets, with higher parasite prevalence in Gabon, Cameroon, and Burkina Faso (Figure 6B, Figure S2D). As expected, STH were largely undetectable in the American dataset (Figure 6B). Notably, and consistent with our previous findings, the number of parasite species was positively associated with Shannon diversity index and Faith’s PD, particularly in the Gabon, Cameroon, and Burkina Faso cohorts (p = 0.000319, p = 1.73E-12, respectively, Tables S4a and S4b, Figure 6C & D).

**Figure 6:**
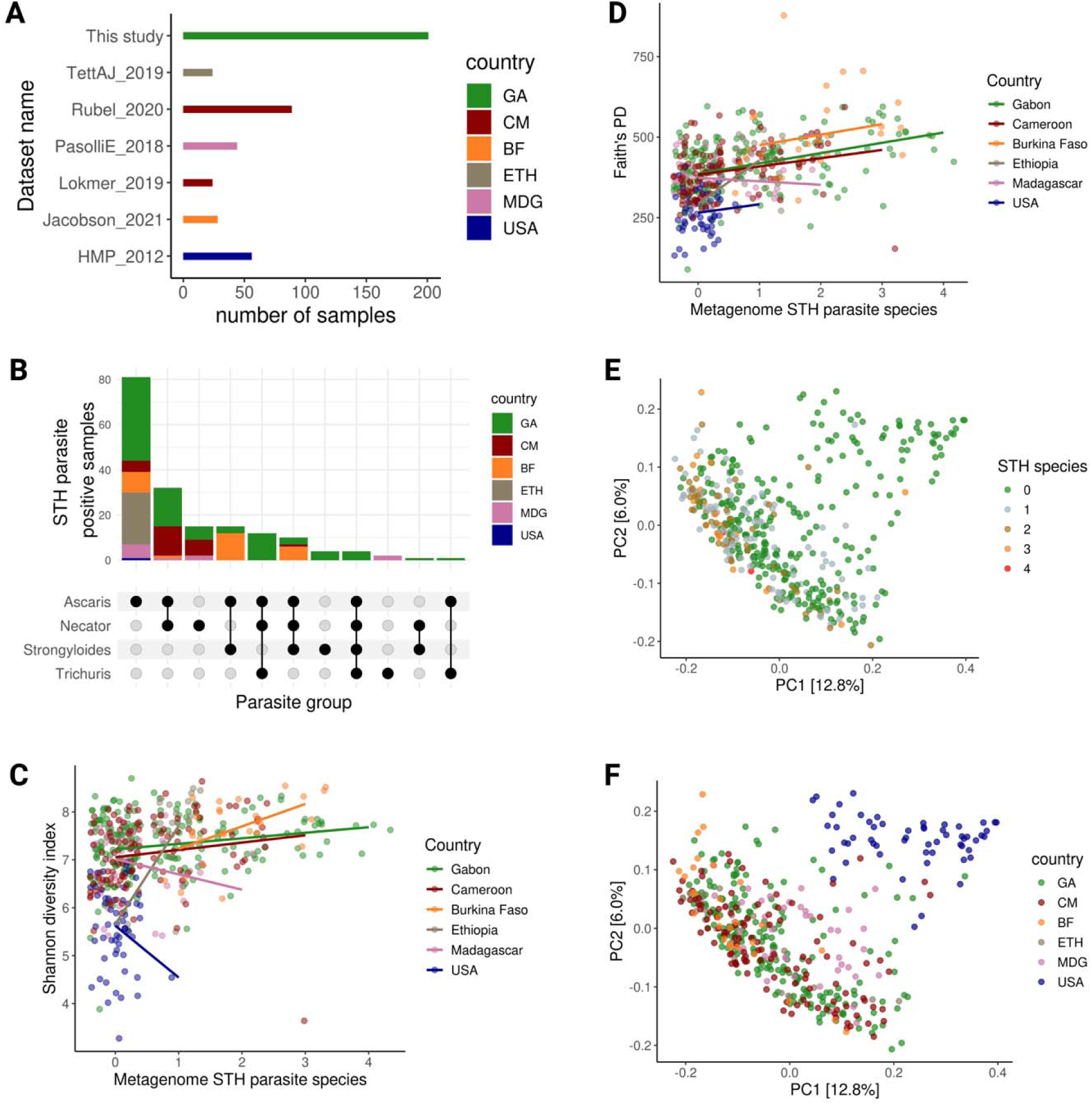
STH parasite richness (the number of different parasite species) significantly contributes to gut microbiome diversity across various countries and independent studies. **A)** The distribution of samples based on the STH parasites detected, categorized by datasets (GA = Gabon, CM = Cameroon, ETH = Ethiopia, GHA = Ghana, MDG = Madagascar, TZA = Tanzania, and USA). **B)** Upset plot showing STH parasites detected from metagenomes across datasets, with colors representing the country of origin. **C)** Correlation between Shannon diversity index and the number of STH parasites across datasets, using a generalized linear model (GLM). **D)** Same as panel C, showing correlation between Faith’s PD and the number of STH parasite counts across datasets (GLM). **E)** Principal Coordinate Analysis (PCoA) of unweighted UniFrac values, with samples colored based on the number of STH parasites detected. **F)** PCoA of unweighted UniFrac values, with samples colored by country.

Weak clustering by the number of STH species was also observed in the PCoA analysis, along with the dataset or country of sampling (Figure 6E & F). PERMANOVA confirmed a significant association between the STH species count and microbiome beta diversity (Bray Curtis, Jaccard, Weighted and Unweighted UniFrac), although the effect sizes were modest (R^2^ < 0.05) (Table S4c). Overall, these results align well with our previous analysis of the Gabon dataset via qPCR and suggest that there is a consistent association between parasite species presence /absence and microbiome diversity across diverse African populations.

To identify specific microbial species consistently associated with the number of parasite species across datasets, we used the overlap of DESeq2 and MaAslin2 results. After filtering for microbial species prevalence, we found 573 and 525 bacteria significantly associated with the number of parasite species using DESeq2 and MaAslin2, respectively (Tables S5a and S5b). Of these, 312 taxa overlapped between the two methods (Figure 7A & B, Table S5c). Using FGSEA, we identified 7 families significantly overrepresented among species positively (n = 4) and negatively (n = 3) associated with the number of parasites (padj < 0.05; Figure 7C, Table S5d&e). Notably, Lachnospiraceae had the highest representation, with 110 species (35.2%) belonging to this family, mostly positively associated with parasite species count. The *Roseburia* and *Agathobacter* genera within Lachnospiraceae showed the highest effect sizes (log2 fold change >1, Figure 7A). Other enriched families included Clostridiaceae, CAG508, and UBA 932. Conversely, Bacteroidaceae had 59 species (19%) mostly negatively associated with the number of parasites. Tannerellaceae and Marinifilaceae were other families enriched in parasite-negative samples. Additionally, Oscillospiraceae and Ruminococcaceae showed positive associations with the number of parasites, although not significant in FGSEA (Figure 7A).

**Figure 7:**
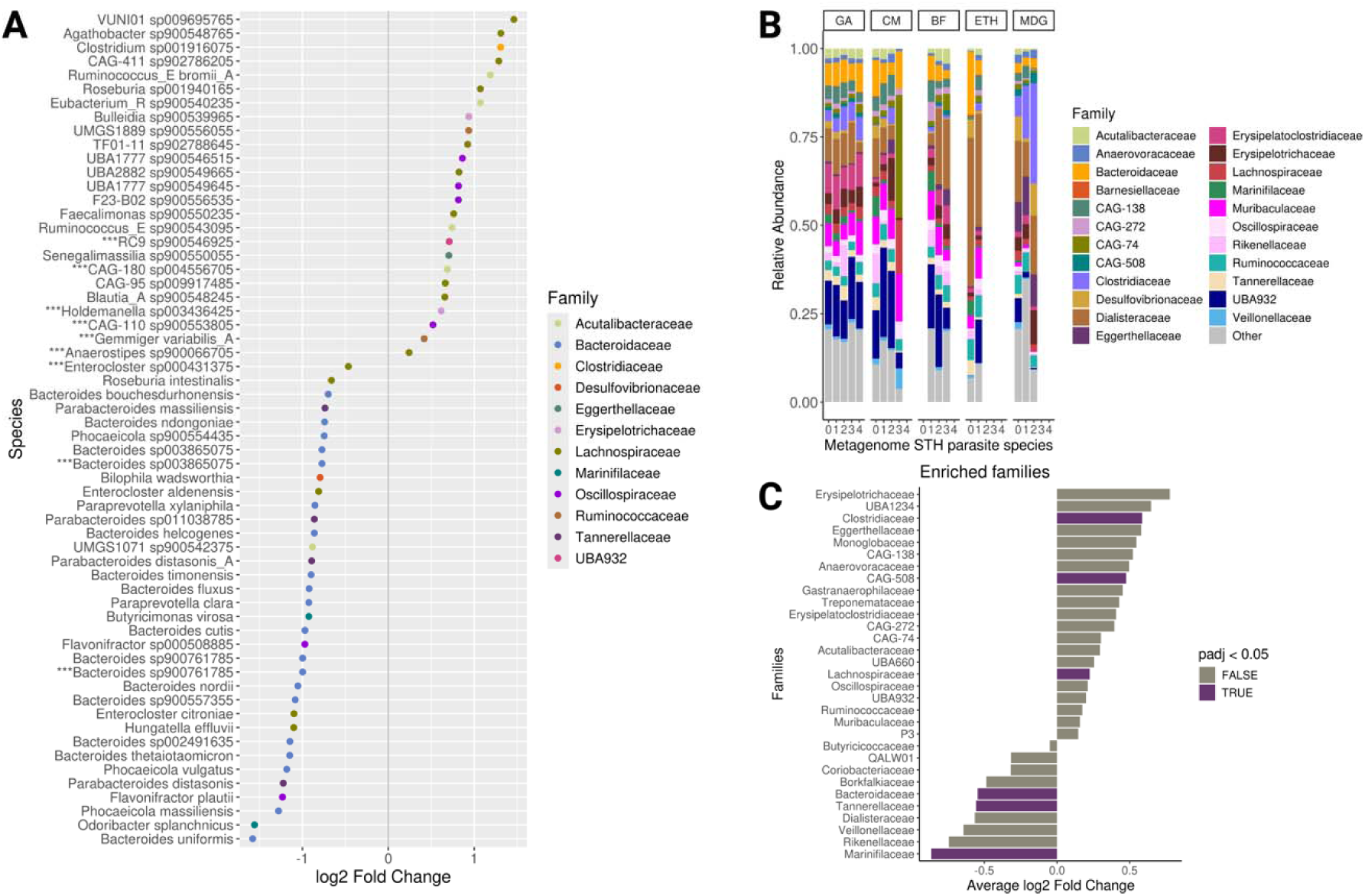
Differential Abundance of Microbial Species by STH Parasite Count Across Multiple Countries and Studies. **A)** Log2 fold changes in the abundance of microbial taxa, as detected by DESeq2 and MaAslin2, based on the number of STH species (1 to 4 STH species or negative = absence of all 4 species). Shown are the top 60 species found by both methods (n = 312). Taxa marked with were also identified in the Gabonese adult female dataset by both methods. Differential abundances were analyzed while accounting for country as a design factor, with multiple inference corrections applied using the Benjamini-Hochberg method and an adjusted p-value threshold of α = 0.05. **B)** Relative abundances of all differentially abundant species identified by both DESeq2 and MaAslin2, colored by family. **C)** Average log2 fold changes of bacterial families significantly enriched in STH parasite-positive or -negative samples, based on Fast Gene Set Enrichment Analysis (FGSEA). Log2 fold changes were derived from DESeq2 analysis.

In our Gabon-only dataset using qPCR for parasite detection, we had previously identified 14 differentially abundant species in adults, and 9 of these species overlapped with the differentially abundant species found in the meta-analysis. This overlap included species from Lachnospiraceae, Bacteroidaceae, Oscillospiraceae, Ruminococcaceae, Acutalibacteraceae, Erysipelotrichaceae, and UBA932 (Figure 7A). Additionally, members of the genus *Roseburia* were positively associated with the number of parasite species in both the adult meta-analysis and in Gabonese children. Together, these findings indicate a consistent signature of parasite infection on the human intestinal microbiome across populations.

Overall, these results indicate that our main conclusions drawn from the Gabonese dataset can be recapitulated using metagenomic detection of parasites and generalized to other metagenomic studies of African adult women. Parasite infection (the number of STH species in a sample) is consistently associated with microbiome richness and with altered abundances of specific microbial taxa across populations in Africa.

## Discussion

Infection with STH is a neglected tropical disease that has a profound yet not fully understood impact on the human gut environment. In this study, we explored the connection between STH parasite infection and the gut microbiota in 155 mother-child pairs from Gabon. Through high-depth metagenomic sequencing and rigorous validation with qPCR, we provide critical insights into how these parasitic infections modulate microbial diversity and community structure in the human gut.

One of the key strengths of our study is the utilization of both qPCR and shotgun metagenomics for detecting STH infections. Traditional microscopy, though effective, is laborious and not routinely performed for fecal microbiome datasets. Our dual approach provided enhanced accuracy in detecting low-intensity infections, with qPCR demonstrating 94% sensitivity. Additionally, our validation of shotgun metagenomics against qPCR revealed robust correlations for most STH species. This methodological advancement not only confirmed the presence or absence of parasites but also highlighted the challenges in accurately quantifying *T. trichiura* due to primer cross-reactivity and complexities in metagenomic read counting.

Our findings indicate a significant positive correlation between the presence of multiple STH parasites and increased alpha diversity, a measure of microbial richness within a sample. This association remained significant even after adjusting for age and sampling location, with a pronounced effect observed in Gabonese children. This result aligns with previous studies from Cameroon, where an increase in alpha diversity was associated with the number of parasites, although no differentiation between children and adults was made ^38^. In contrast, a study conducted in Tanzania reported an increase in alpha diversity among parasite-infected mothers, with only bacterial richness increasing in infected children ^43^. These discrepancies may be attributed to regional differences in parasite species and host responses.

Our meta-analysis of additional African metagenomic datasets confirmed the consistent association between parasite species richness and gut microbiome diversity across diverse populations. This broader perspective enhances the generalizability of our findings and underscores the utility of metagenomic approaches in epidemiological studies of parasite-microbiota interactions. The observed positive correlation between parasite species richness and gut microbiome alpha diversity, particularly pronounced in children but evident across all age groups, suggests a universal pattern of microbial community response to STH infections. This aligns with global reports indicating increased microbial richness in helminth-infected individuals ^44,45^, suggesting potential mechanisms such as parasite-induced immune modulation or dietary changes that promote microbial diversity in the gut.

We observed specific microbial taxa associated with parasite infections. Notably, species from the Bacteroidaceae and Lachnospiraceae families showed negative associations with parasites, in both Gabonese adults and the meta-analysis dataset. Additionally, *Bifidobacterium* demonstrated a strong negative association with parasites in Gabonese children. Conversely, certain Bacteroidales species exhibited a positive association with parasites across all datasets. Previous studies in humans have reported an increase in Bacteroidales in parasite-infected samples ^38,46^, while mouse models of helminth infection have shown a decrease in Bacteroidales ^18^. These results suggest that Bacteroidales may respond differently to parasites, indicating that contradictory outcomes in previous studies could be attributed to insufficient taxonomic resolution. Previous reports have also shown an increase in the abundance of Clostridiales in the presence of STH parasites, accompanied by a shift from Bacteroidota to Firmicutes ^43,47–49^. It is important to note that various species within these families associate differently with parasite infections, indicating the complexity of STH-parasite-microbiota associations. Interestingly, several of the bacterial taxa that correlated positively with parasite species count in our data and meta-analysis are known mucus degraders ^50^. Studies on mice have demonstrated that the absence of mucin production impairs their resistance to infection by *Trichuris muris*, a common mouse parasite ^51^. It is plausible that when STH parasites colonize the gut, the abundance of certain bacterial mucin degraders diminishes due to competition for mucin resources. Simultaneously, the parasites may promote the growth of other bacteria capable of mucin degradation.

The co-occurrence network analysis provided further insights into the ecological relationships between STH parasites and the gut microbiota. We observed significant associations between specific bacterial species and STH parasites, particularly in adults where all four STH species showed distinct co-occurrences with microbial taxa. Ascaris exhibited the highest number of unique co-occurrences, often with Firmicutes and Bacteroidota, indicating potential niche preferences and microbial community adaptations in response to parasitic colonization. These results suggest that while STH parasites influence specific microbial niches, they do not induce a uniform shift in microbiome composition, highlighting the nuanced nature of parasite-microbiota interactions in the gut ecosystem.

Despite the strengths of our study, such as high-depth metagenomic sequencing and robust validation techniques, there are limitations that warrant consideration. The uneven distribution of parasite-infected individuals across different sites and age groups within the Gabonese cohort may have introduced bias in our results. Additionally, we did not account for potential confounding factors such as socioeconomic status, dietary habits, or medication use, which could independently influence microbiome composition. Future research should incorporate comprehensive demographic and environmental data to better understand the multifaceted influences on parasite-microbiota associations.

In summary, our study presents the first assessment of correlations between STH parasite infections and gut microbiota associations in rural communities in Gabon. By including stool samples from villages in a remote community in southern Gabon, we expand the epidemiological understanding of STH and their associations with the microbiome in these populations. Furthermore, our research enhances our comprehension of parasite-microbiome associations across diverse populations in Africa. Overall, our findings underscore the significance of parasite-microbiota associations, particularly in children who are commonly targeted for control interventions through mass drug administration (MDA). As these associations may persist throughout life and impact host health, they should be taken into account when optimizing deworming programs in areas endemic to STH parasites.

Our findings have significant implications for public health strategies, particularly in the design of deworming programs targeting vulnerable populations, such as children in endemic areas. By elucidating the association of STH infections with gut microbiome diversity and community structure, our study highlights the potential role of microbiota modulation in enhancing host resilience against parasitic infections. Further research is needed to explore the mechanistic underpinnings of parasite-microbiota interactions and their implications for host health and disease susceptibility in diverse epidemiological contexts.

## Conclusion

In conclusion, our study advances the understanding of parasite-microbiota interactions in African populations by integrating high-resolution metagenomic analyses with robust validation techniques. By demonstrating consistent associations between STH infections and gut microbiome diversity across diverse populations, we provide insights into the ecological dynamics of parasitic infections and their potential implications for host health. Moving forward, continued research efforts should focus on elucidating the mechanistic pathways linking parasite colonization to microbiome modulation and exploring novel therapeutic strategies leveraging microbiota-targeted interventions for improving health outcomes in parasite-endemic regions.

## Materials and Methods

### Study Populations

The study was conducted at the Centre de Recherches Médicales de Lambaréné (CERMEL), Gabon between November 2017 and February 2018. Study participants were mother and child pairs from multiple ethnic groups living in Lambaréné in the vicinity of CERMEL and those living in 5 Ikobey villages including; villages surrounding Tchibanga, Tranquille, Soga, Erouba, and Ikobey of the Southern part of the country (Figure 1A). Given their similar lifestyles, all participants from the 5 villages are grouped as one community and labeled "Ikobey’’. Their subsistence practices are centered around hunting, agricultural crop cultivation, and livestock rearing ^52,53^. In contrast, participants from and within the vicinity of Lambaréné, a semi-urban town in the Moyen-Ogooue province of Gabon, mostly practice fishing, farming, and a semi-westernized lifestyle. The metadata associated with samples in this study is shown in Table S6.

### Approval and Informed consent

The study was reviewed and approved by the "Comité National d’Ethique ’’ of Gabon with registration number PROT N°0025/2017/SG/CNE. Stool samples were collected after participants provided written informed consent or assent. Sample collection - Stool samples were collected in sterile plastic containers from both mothers and their children. Stool samples collected on-site (CERMEL) were aliquoted and stored at -20 °C within 2 hours of sample collection. Samples collected in the field (Ikobey) were transported in cold boxes to CERMEL and later stored at -20 °C. Samples were later transported to Germany on dry ice where they were stored at -80 °C before DNA extraction, qPCR, and sequencing. Publicly available datasets - We utilized publicly available African gut metagenome datasets for studies performed in Cameroon ^25,38^, Burkina Faso ^39^, Ethiopia ^40^, and Madagascar ^41^. The two Cameroon cohorts were sampled from ethnically diverse populations that practiced pastoralist, agropastoralist, or hunter-gatherer lifestyles ^38^, or from hunter-gatherers, farmers, or fishing populations in Southwest Cameroon ^25^. The Burkina Faso cohort comprised samples collected from healthy volunteers in a single village in Burkina Faso ^39^. The Madagascar cohort comprised samples collected from two ethnic groups in a remote rainforest region of north-eastern Madagascar ^41^ and the Ethiopia cohort included samples from healthy adult females ^40^. We also included one USA cohort, which is a subset of the Human Microbiome Project (HMP) dataset ^42^. Only female subjects who were described as healthy (mostly self-report) and ≥18 years old were included. The combined dataset used for our meta-analysis comprised 466 metagenome samples. The metagenomes from the 5 African and one USA cohorts were sequenced on the Illumina platform at an average depth of 45 M read pairs. These metagenomes were retrieved from Sequence Read Archive (SRA) and processed as described here ^54^ and here ^38^. The USA cohort was not included in the statistical analysis and was used for illustrative purposes only. The metadata associated with samples in the public datasets is shown in Table S7 and Table S8.

### Detection of parasites by microscopy

Stool samples collected on-site (Lambaréné) were examined for the presence of parasites by using the Kato-Katz ^55^ concentration technique to detect and quantify parasite eggs. We used a modified Harada-Mori ^56^ technique to detect the larvae of *N. americanus* and *S. stercoralis*. Samples were visualized independently by two laboratory technicians under the standard light microscope to identify and count the number of parasitic elements (eggs/larvae). The average of the two readings was recorded, and final egg counts were converted to eggs per gram of feces using standard CERMEL parasitology laboratory protocols for reporting egg counts ^28^. Microscopy analysis was not performed on field (Ikobey) samples due to logistic reasons. To obtain a comprehensive picture of parasite prevalence in these samples, qPCR targeting all four parasites mentioned above was performed on all samples, including samples from Lambaréné for which microscopy results were available (see "Detection of parasites by quantitative PCR" below). Microscopy-based detection on Lambaréné samples was used to provide support to the qPCR data. To evaluate the sensitivity and specificity of both methods, we considered as a true positive any sample that tested positive by either qPCR or microscopy and a true negative any sample that tested negative by either method.

### Detection of parasites by quantitative PCR

Diagnosis of STH parasites was performed by multiplex qPCR using protocols as described elsewhere ^57,58^. The target species were *A. lumbricoides*, *T. trichiura*, *S. Stercoralis*, and the hookworm species *N. americanus*. The primers, probes, and reaction conditions for amplification were based on published protocols ^58,59^ with minor modifications. Primer sequences are given in Table S10. An internal control DNA from Phocine Herpesvirus 1 (PhHV-1) was added into the master mix to control for DNA amplification inhibition. The PCRs were split into two multiplex assays combining *A. lumbricoides, T. trichiura*, and PhHV-1 as targets (assay 1) and *S. stercoralis, N. americanus*, and PhHV-1 as targets (assay 2). The positive controls were plasmids carrying the targeted regions of the four target parasites, kindly provided by the Institute of Tropical Medicine (University of Tübingen). The target genes inserted in these plasmids were an 87 -bp fragment of the ITS1 sequence of *A. lumbricoides*, a 101-bp fragment of the 18S rRNA gene sequences of *S. stercoralis* and *T. trichiura,* and a 101-bp fragment of the ITS2 gene sequence of *N. americanus.* Wells without any template served as negative controls, and all samples, including controls, were tested in duplicate. The total reaction volume was 12.5 μl, which included: 6.25 μl Qiagen HotStarTaq Master Mix (QIAGEN GmbH, Hilden, Germany), 1 μl of 25 mM MgCl_2_, 0.25 μl of 5 mg/ml BSA, 0.1 μl species-specific primers (final concentration of 800 nM), 0.0625 μl of probes (final concentration of 500 nM), 1.5 μl of PhHV-1 plasmid DNA (diluted to 10^-7^), 2 μl of template DNA, 0.2125 μl of PCR-grade water (Sigma-Aldrich, Darmstadt, Germany). The qPCRs were conducted for 40 cycles using a Bio-Rad CFX384 Touch Real-Time PCR Detection System (Bio-Rad Laboratories, Germany). Only samples with cycle threshold (Ct) values ≤ 35 were considered positive. All negative controls hat Ct values above 35.

DNA extraction and shotgun metagenomic sequencing - Total DNA from fecal samples was extracted from approximately 250 mg of frozen stool aliquots. DNA extractions were performed using the Qiagen DNeasy PowerSoil HTP 96 kit (QIAGEN GmbH, Hilden, Germany) following the manufacturer’s instructions. Extraction blanks were included to control for reagents and environmental contamination. Extractions were performed under sterile conditions in a laminar flow hood, and the eluted DNA was stored at -20 °C following quantification with Qubit 4 fluorometer (Fisher Scientific GmbH, Schwerte, Germany). Genomic DNA libraries were constructed using the Nextera protocol ^60^ with in-house produced Tn5 enzyme ^61^. Briefly, DNA was diluted to 5 ng/µl and tagmented with the Tn5 transposase, followed by amplification using a combination of paired-barcoded primers for 12 cycles. Libraries were pooled and quantified with a Qubit, followed by size selection for the 300-700 bp range on a BluePippin. The resulting libraries were then sequenced on an Illumina HiSeq3000 at the Max Planck Institute for Biology in Tübingen.

### Processing of sequences and taxonomic profiling

All Illumina shotgun raw sequences were processed similarly to Youngblut and colleagues ^54^. Briefly, raw reads were validated with fqtools v.2.0 ^62^ and de-duplicated with the "clumpify" command from bbtools v37.78 (https://jgi.doe.gov/data-and-tools/bbtools/). Adapters were trimmed and quality controlled with Skewer v0.2.2 ^63^, and the "bbduk" command from bbtools. Human genome reads mapped to the hg19 assembly were filtered with the "bbmap" command from bbtools.

Quality control (QC) reports for all reads were generated with fastqc v0.11.7 (https://github.com/s-andrews/FastQC) and multiQC v.1.5a ^64^. The filtered reads were then subsampled to 5 million reads per sample, and a combination of Kraken 2 ^65^ and Bracken v2.2 ^66^ was used to classify filtered, quality-controlled reads with a custom database build from Genome Taxonomy Database (GTDB, Release 202; https://gtdb.ecogenomic.org/). For parasites profiling from the filtered reads using KrakenUniq ^36^, we first used Kraken2 to separate reads that were classified to bacteria/archaea, then used the remaining unclassified reads to map to our custom-built reference genome database of the 4 STH parasites ^32^. We included 4 STH parasites: *A. lumbricoides*, *S. stercoralis*, *N. americanus*, and *T. trichiura* in the database. Bacteria and Archaea classification with Kraken2 were resolved to species. KrakenUniq classification of STH parasites was resolved with unique k-mers. K-mer counts <1000 were considered as noise and replaced with zeros as suggested by the authors of KrakenUniq ^36^. Abundances (k-mer counts) were converted to presence/absence.

### Assessing parasite genome assembly quality

All 4 STH parasite genomes used for profiling with KrakenUniq were downloaded from NCBI. Accession numbers are provided in Table S9. Metagenome read coverage across the entire length of each STH parasite genome was assessed with Bowtie2 ^67^. The Bowtie2 index of the 4 reference genomes was built with bowtie2-build. Nucleotide coverage across each length of the contig was analyzed in R ^68^. BLASTn searches on contigs with the highest coverage revealed hits to target parasite species for all 4 STH parasites. The sample processing workflow is shown in Figure S3.

### Statistical Analysis

The mean ages and proportions of STH parasite infections were compared between groups using the Wilcoxon test. All diversity measures were obtained from shotgun metagenomic data. Reads were randomly subsampled to 5000000. Diversity was assessed using QIIME 2 ^69^ at a rarefaction depth of 300000 reads. The alpha diversity measures assessed include Shannon’s index (which takes into account the abundances and evenness of species), and Faith’s PD (which takes into account the phylogenetic relatedness of species). Beta diversity was assessed using Bray Curtis (takes into account species abundances) and Jaccard (measures presence/absence of species) dissimilarity indices. For the Gabon-only cohort, metadata variables tested with alpha and beta diversity in R include the number of parasites, age group, and location. Linear models were used to assess the relationships between alpha diversity and the number of parasites while accounting for age group and location. PERMANOVA test was performed in R using the adonis function of the vegan package ^70^. PERMANOVA test was done on Bray Curtis, Jaccard, Weighted, and Unweighted UniFrac distances to assess parasite-microbiota associations while accounting for age group and sampling location differences. Similar measures of alpha and beta diversity were employed in the meta-analysis. Linear mixed models were used to assess the relationship between the number of parasites and alpha diversity while controlling for dataset ID (random effect). PERMANOVA was also performed on beta diversity measures with 999 permutations constrained by dataset ID.

### Differential abundance analysis was performed using DESeq2 ^33^ and MaAslin2 ^34^

While using DESeq2, geometric means were calculated before estimating size factors, and apeglm ^71^ was used for LFC shrinkage. For MaAslin2, species counts were log-transformed and normalized by total sum scaling (TSS). Minimum abundance and minimum prevalence were both set at 0.0, and a linear model was chosen for the analysis method in MaAslin2. For both methods, the fixed effects were the number of parasites and location (Gabon-only cohort) or number of parasites, country, and age (meta-analysis) in the design factor. The age and number of parasites (continuous variables) were standardized by applying z-scores in MaAslin2. Microbiome input data were unnormalized sequence counts after single reads were removed to avoid sequencing errors. A further prevalence filtering (>80%) was performed on both separate Gabon mothers/children and the meta-analysis datasets before analysis with DESeq2 and MaAslin2. The multiple inference correction used is Benjamini-Hochberg. The adjusted p-value was set at (alpha = 0.05) and entries with an NA value were removed. Enriched families were assessed with the R package *fgsea* ^72^ for GSEA ^35^. For this analysis, all the DA species (not only significant species at p < 0.05) from DESeq2 and MaAslin2 were separately grouped into families. For each species, their corresponding log2 fold change was used for ranking. The minimum set size (number of DA species in a Family) was set as 2 for both Gabon only and meta-analysis datasets, and the maximum set size was 500 and 800, respectively. This means microbial families with less than 2 DA species (minSize) and those with more than 500 or 800 DA species (maxSize) are excluded from the analysis. The maxSize values are different between the two datasets, as more DA species were observed in the meta-analysis dataset. For estimating p-values more accurately, the eps argument of *fgsea* was set to zero.

### Co-occurrence abundance analysis was performed with the R package cooccur ^73^

First, the microbiota species abundance counts were split between adults and children. Species counts were then transformed to relative abundances, filtered to a >50% prevalence, and then converted to presence/absence (1/0) in separate datasets. Similarly, the number of parasites from KrakenUniq was converted to presence/absence and then joined to the microbiota presence/absence table in the separate datasets. Probabilities of co-occurrence between microbiota and STH parasites were defined as negative if p_lt < 0.05, or positive if p_gt < 0.05, where p_lt and p_gt are probabilities that two species do not co-occur less or more frequently than expected. Only the species with significant probabilities of cooccurrences with the 4 STH parasites were included in the network graphs. The weights are the differences between the observed and expected frequencies of co-occurrence normalized by the sample size (n=155) for adults and children respectively.

The following R ^68^ packages were used for data processing and visualizations: phyloseq ^74^, microbiome ^75^, dplyr ^76^, tidytable ^77^, tidygraph ^78^, broom ^79^, igraph ^80^, ggplot2 ^81^, ggraph ^82^, UpsetR ^83^, complexUpset ^84^, fgsea ^72^, MaAsLin2 ^34^, concur ^73^

## Supporting information

All Supplemental Material

## Acknowledgments

We are grateful to the laboratory personnel of CERMEL-Gabon and members of the Department of Microbiome Science at the MPI for Biology, in particular Silke Dauser, for technical assistance. This work was funded by the Max Planck Society.

## Data availability statement

The authors confirm that all data supporting the findings of this study are included within the article and its supplementary materials. The raw sequence data for the Gabon cohort can be accessed from the European Nucleotide Archive under the study accession number

PRJEB46788. Additionally, the meta-analysis dataset and references are provided in the supplementary tables.

## Appendices

List of tables

List of figures

Table 1

Supplementary tables

## Supplementary Figures

**Figure S1:**
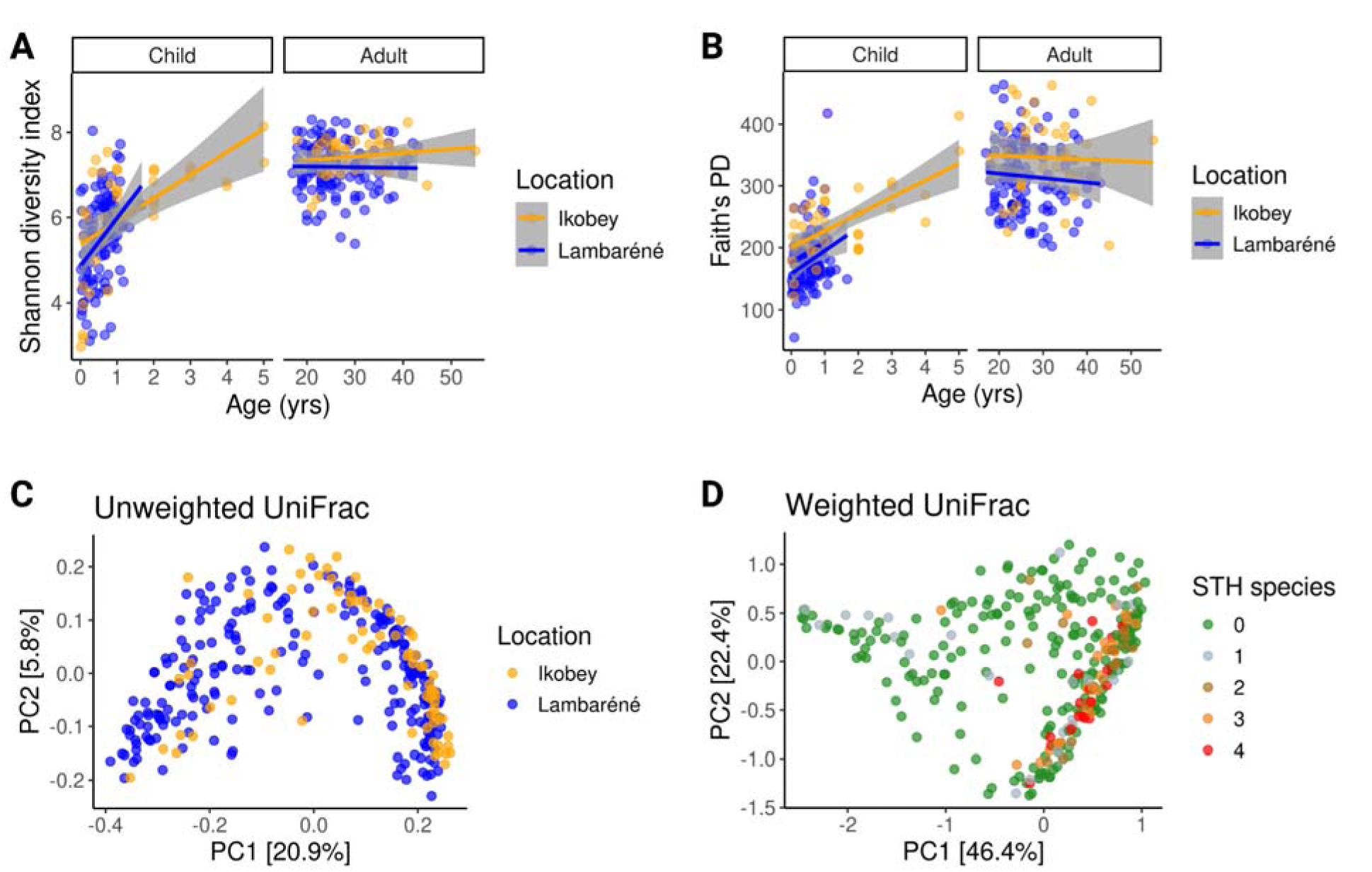
Gut Microbiome Diversity by Age Group and Location. **A)** Correlation of the gut microbiome Shannon diversity index with age and location (correlation method: GLM). For children, *R^2^* = 0.33 in Ikobey and *R^2^* = 0.15 in Lambaréné; for adults, *R^2^* = 0.02 in Ikobey and *R^2^* < 0.01 in Lambaréné **B)** Correlation of Faith’s Phylogenetic Diversity (PD) with age and location (correlation method: GLM). For children, *R^2^* = 0.45 in Ikobey and *R^2^* = 0.08 in Lambaréné; for adults, *R^2^* < 0.0 in both Ikobey and Lambaréné. **C)** Principal Coordinate Analysis (PCoA) of unweighted UniFrac distances among all samples (n = 310), with samples colored by location. **D)** PCoA of weighted UniFrac distances among all samples, with samples colored by STH species counts.

**Figure S2:**
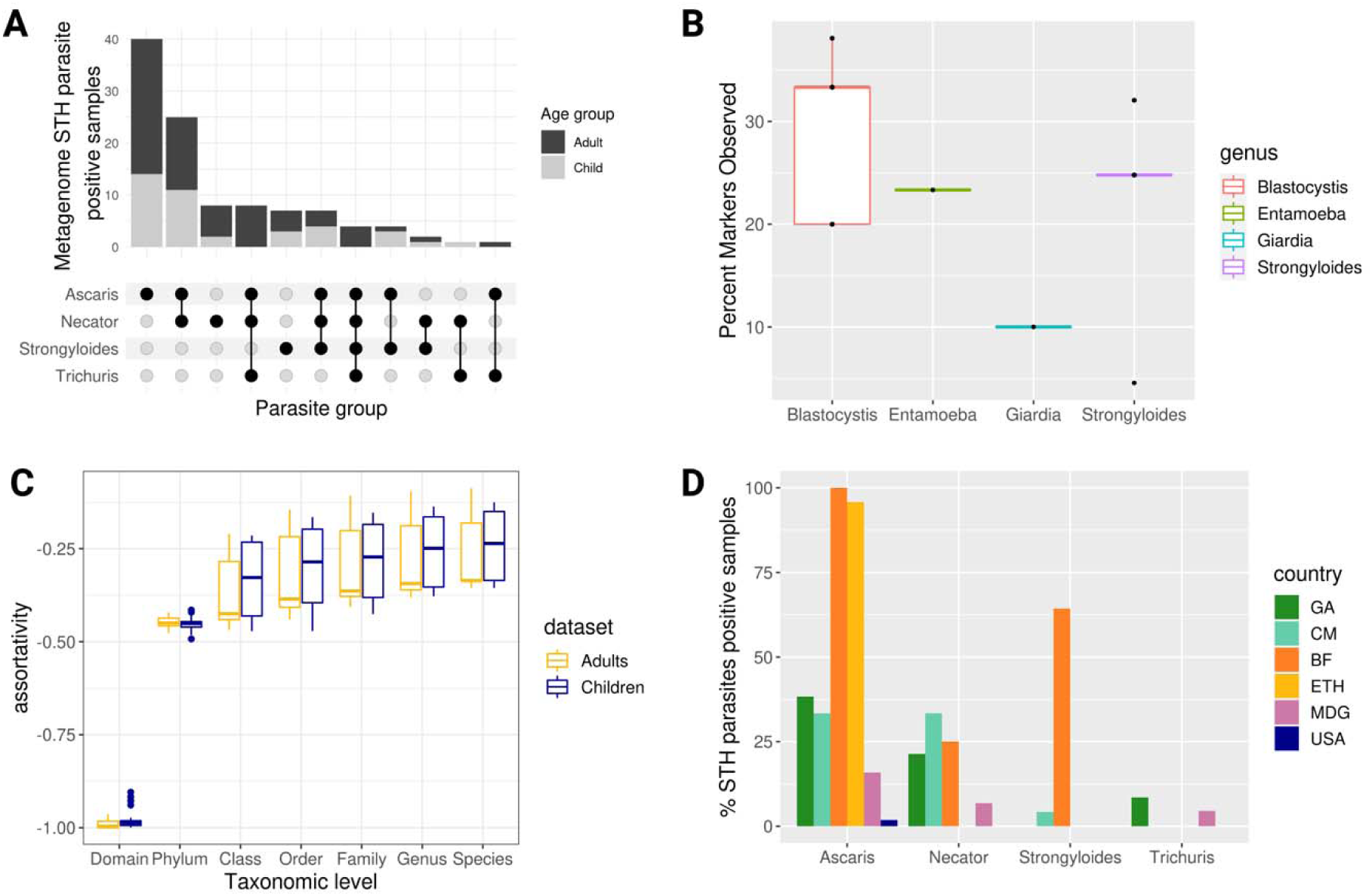
Detection of soil-transmitted helminths (STHs) in gut metagenomes of African adult females. **A)** Upset plots showing the detection of STH parasites in metagenomes from Gabonese mother-child pairs. **B)** Percentage of parasite markers observed in Gabonese mother-child metagenomes using the Eukdetect database. **C)** Boxplots of assortativity coefficients (node connections) at different taxonomic levels for the graphs in panel A. For each graph, a random subset of 100 replicates with 230 nodes was analyzed, and assortativity values were calculated for each taxonomy level at each iteration. **D)** Bar plots showing the percentage of STH-positive samples detected from metagenome across different datasets, colored by country

**Figure S3:**
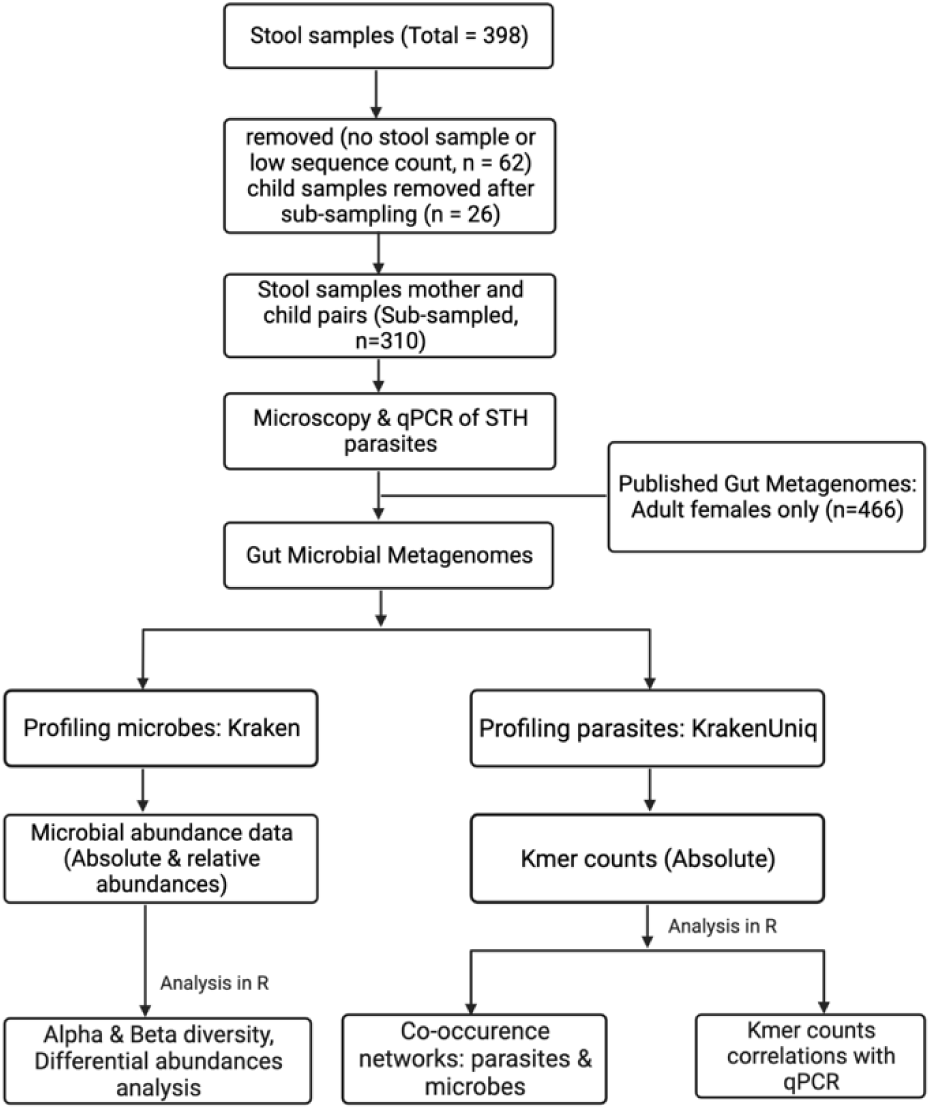
Sample Processing Workflow. Participants without stool samples and samples with low sequencing counts (<100,000 reads) were excluded from downstream analysis (n = 62). An additional 26 samples were removed to focus on mother-child pairs. Publicly available metagenomes from adult women in studies conducted across four African countries (n = 265) and this study’s cohort (n = 201) were included in the meta-analysis. Microbial abundance was estimated using Kraken, and KrakenUniq was used to estimate the number of unique k-mers assigned to each STH parasite.

